# The structure of the humanised A33 Fab C226S variant, an immunotherapy candidate for colorectal cancer

**DOI:** 10.1101/2022.06.21.497004

**Authors:** Jiazhi Tang, Cheng Zhang, Paul Dalby, Frank Kozielski

## Abstract

Colorectal cancer (CRC) causes the second highest cancer-related deaths worldwide. The human A33 antigen is a validated immunotherapy target, which is homogeneously expressed in 95% cases of primary and metastatic colorectal cancers. In this article, we report the structure of a humanised antigen-binding fragment A33 (A33 Fab), a therapeutic antibody candidate, in two different crystal forms. Insights into the structural features of A33 Fab are provided with a focus on the ‘grafted’ complementarity-determining regions (CDRs) and the switch linker between the variable and the constant regions.

**Synopsis:** The crystal structure of humanised A33 Fab, targeting colorectal cancer related antigen, was determined in two different space groups.

## 1. Introduction

Colorectal cancer (CRC) is a common cancer type worldwide with a high metastases rate and which causes the second most cancer-related deaths (Bray, Ferlay et al. 2018). It is reported that approximately 20% of newly diagnosed patients are identified with metastases in liver, lungs, lymph nodes, peritoneum or soft tissues (Riihimäki, Hemminki et al. 2016). Recently, this percentage dropped significantly owing to the screening methods including fecal occult blood tests, colonoscopy and colonography, which increase the possibility to diagnose CRC at earlier stages (Noguchi, Ritter et al. 2013). However, the disease control for patients with advanced-stage CRC remains to be a challenge and intensive therapies including irinotecan or oxaliplatin-based chemotherapies, signalling inhibitors and antibodies are required (Van Loon and Venook 2011). Though immunotherapy featured by therapeutic antibodies has been widely accepted for tumour treatment, currently no approved immunotherapeutic antibody is available clinically against CRC.

Human A33 antigen, a *M*_r_ 43,000 glycoprotein, has been selected as an immunotherapy target. The expression of the A33 antigen is restricted to the epithelia of the lower gastrointestinal tract, as well as to carcinoma lesions that originate from the rectal and colonic mucosa (Welt, Ritter et al. 2003). It has been reported that the A33 antigen expressed in metastatic colorectal cancers share 95% similarity (GarinChesa, Sakamoto et al. 1996).

The original monoclonal antibody (mAb) against A33 originating from murine IgG2a has undergone a set of preclinical analysis followed by a series of Phase I clinical trials. Haematological toxicity was reported as the major limiting toxicity which is triggered by the human anti-mouse antibody (HAMA) response (King, Antoniw et al. 1995). To lower the toxicity as well as to extend the half-life of mAb A33, a chimeric A33 antibody was developed by combining the variable region of mAb A33, which is responsible for antigen binding, with the constant region of the human antibody LAY. The chimeric A33 antibody has 75% sequence identity to human IgG1 (King, Antoniw et al. 1995). Though the human immune system reacted with a mild response to chimeric A33, the human anti-chimeric antibody (HACA) response is still triggered. The failure of the chimeric antibody implied further humanization of chimeric A33 antibody was needed to lower the immune response. Thus, a fully humanized A33 Fab was developed by grafting only the CDRs from mAb, which are responsible for antigen binding, into the variable region framework of the human antibody (King, Adair et al. 2001).

Several A33 Fab variants have been developed by Union Chimique Belge Pharma (UCB) for the treatment of colorectal cancer. These were tested as therapeutic antibodies either in their isolated forms, or chemically cross-linked using trimaleimide, which produces a trivalent Fab fragment that binds antigen with greater avidity, to further increase drugability (King, Antoniw et al. 1995). However, none of these particular variants progressed into the market, either due to toxicity or stability problems. Though several manuscripts have been published on the conformational flexibility and kinetic stability of the A33 Fab mutant H/C226S (named as A33 Fab throughout this article) (Chakroun, Hilton et al. 2016, Zhang, Samad et al. 2018), no crystal structure of any of the A33 Fab variants is available up till now. Here we report the structure of the fully humanised A33 Fab H/C226S. In the H/C226S variant the cysteine is mutated to serine at the 226^th^ residue of the heavy chain to eliminate intermolecular dimerisation (Chakroun, Hilton et al. 2016).

We crystallized the A33 Fab in two different space groups, *P1* and *P6_5_*, and solved both structures to 2.5 and 2.2 Å resolution, respectively. We provide insights into the general structure of the A33 Fab as well as the six CDRs that originated from the A33 mAb. Meanwhile, we comparatively analysed the A33 Fab with Certolizumab, a humanised therapeutic antibody developed from the same humanization framework but targeting a different antigen and elucidated the differences in their CDR- and switch regions.

To evaluate to what extent the crystal structures in space groups *P1* and *P6_5_* could reflect the *in vitro* protein dynamics and stability, the *in silico* ΔΔ*G* was calculated for various mutations previously studied *in vitro* (SI), based on each of the two structures (Zhang, Samad et al. 2018). In addition, *in silico* modelling was carried out to predict possible conformations of the two loop regions missing in the x-ray structures.

Finally, we used molecular dynamics simulations to evaluate the dynamics and range of conformations accessible to A33 Fab, particularly through variability in the elbow angles as a result of conformational flexibility in the switch regions.

## 2. Materials and methods

### 2.1 A33 Fab sample preparation

Expression, purification and buffer exchange of A33 Fab has been described previously (Zhang, Samad et al. 2018). Purified A33 Fab was further concentrated to 22.2 mg/ml in water with a 30 kDa Vivaspins (Generon Ltd., Bershire, UK). Prior to crystallization, samples were diluted with PIPES buffer (10x stock of 50 mM PIPES pH 7.0 and 100 mM NaCl) and MilliQ water to a final concentration of 10 mg/ml, 15 mg/ml and 20 mg/ml, respectively.

### 2.2 Protein crystallization

Purified A33 Fab was used to screen for crystallization conditions using the Low Ionic Strength Screen Kit (Hampton Research). Hanging drops were set up in 24-well Linbro plates (Molecular Dimensions) with 4 μl protein solution, 2 μl buffer reagent and 5 μl precipitant reagent, which were equilibrated against 1 mL of 24% w/v PEG3350 reservoir as described in the Low Ionic Strength Screen handbook (Hampton Research). Drops were set up at both 277 K and 291 K.

Crystals grew in two different space groups. Triclinic crystals (Space group *P1*, 2.5 Å resolution) grew from 4 μl protein solution at 20 mg/ml mixed with 2 μl 50 mM Glycine pH 9.0 and 5 μl 16% w/v PEG3350 at 291 K. The crystals were cryoprotected with 60 mM Glycine pH 9.0, 19% PEG3350 and 15% Glycerol and flash-frozen in liquid nitrogen. Hexagonal crystals (Space group *P*6_5_, 2.2 Å resolution) grew from 4 μl protein solution at 20 mg/ml mixed with 2 μl 50 mM Citric Acid pH 4.0 and 5 μl 8% w/v PEG3350 at 291 K. The crystals were cryoprotected with 60 mM Citric Acid pH 4.0, 10% PEG3350 and 15% Glycerol and flash-frozen in liquid nitrogen.

### 2.3 Data processing and structure determination

X-ray diffraction data were collected on Diamond beamline I04-1 for the hexagonal and I24 for the triclinic structure (Diamond Light Source, Harwell, UK). Data were indexed and integrated with *iMOSFLM* (Battye, Kontogiannis et al. 2011). Data reduction and scaling was accomplished using *SCALA* (*Evans 2006*) within the *CCP4* suite of programs (Collaborative 1994).

The first structure, which is in triclinic form, was determined using *phenix.phaser* (Liebschner, Afonine et al. 2019). A chimeric model was applied as the probe for molecular replacement consisting of the heavy chain of the humanized RK35 antibody (Apgar, Mader et al. 2016) and the light chain of anti-ErbB2 Fab2C4 (Vajdos, Adams et al. 2002). The second structure in hexagonal form was then solved based on a partially refined model obtained from the triclinic crystals. Refinements were carried out by *phenix.refme* (Liebschner, Afonine et al. 2019) iterated with real-space manual rebuild using *Coot* (Emsley, Lohkamp et al. 2010). The first rounds of refinements were done in *phenix.refine* with rigid body and update water functions applied. Once a sharp decrease in *R_free_* value had been observed, simulated annealing and translation–libration–screw (TLS) parameterization were included separately for systematic comparison. Refined models were manually rebuilt in *Coot* and iteratively refined by *phenix.refine*.

### 2.4 Prediction of mutational ΔΔ *G* by Rosetta *cartesian_ddg*

Single point mutations were evaluated in silico using the Rosetta *cartesian_ddg* (Park, Bradley et al. 2016) method for the variants previously studied in vitro (Zhang, Samad et al. 2018). For each mutation, a mutfile was supplied to specify the target mutation. Average ΔΔ *G* values were calculated based on three iterations for both the wild type and mutant, and were correlated with in vitro *T_m_* and aggregation kinetics, *ln*(*v*). Calculations were submitted to the UCL Myriad High-Performance Computing Facility (Myriad@UCL) with Rosetta Version 2018.48.60516-mpi.

### 2.5 Homology modelling

*RosettaCM* (Song, DiMaio et al. 2013) was used to perform homology modelling by applying the Fab A33 full-length amino acid sequences (SI) to the raw crystal structures of P1 and P65. For each homology modelling, more than 20,000 structures were generated after the *RosettaCM Hybridize* step. The top structure with all the five disulfide bonds was selected for the subsequent *RosettaCM Final relax* step. In the *Final relax* step, more than 20,000 structures were obtained and the top structure with all the five disulfide bonds was selected. Jobs were submitted to the UCL Myriad High-Performance Computing Facility (Myriad@UCL) with Rosetta Version 2018.48.60516-mpi.

### 2.6 Molecular dynamics

The Gromacs (Pronk, Páll et al. 2013, Abraham, Murtola et al. 2015) software was used to perform molecular dynamics, using the full-residue structures modelled based on *P1* and *P*6_5_crystals. The PDB2PQR (Dolinsky, Nielsen et al. 2004) webserver was used to determine the protonation states of chargeable residues at pH 7. The simulation was carried out under *OPLS-AA/L* all-atom force field at 300 K and 1 bar. The protein molecule was placed within a cubic box with 1 nm distance between the protein and the box. The box was solvated with *SPC/E* waters, and Na^+^ and Cl^-^ ions to neutralise and provide ionic strength at 50 mM to the system. The system was then energy minimised and equilibrated around the solute protein, under an NVT (constant Number of particles, Volume, and Temperature) and an NPT (constant Number of particles, Pressure, and Temperature) ensemble. Production run was finally conducted for 100 ns. Six repeats were performed for each condition. Jobs were submitted to the UCL Myriad High-Performance Computing Facility (Myriad@UCL) using gromacs/2019.3/intel-2018.

### 2.7 Elbow angle calculation

The elbow angle of the Fab is the angle between the pseudo-twofold axes between the variable and constant domains. It is calculated using the Pymol (Version 2.3.2) elbow_angle.py script, setting the V_L_ domain as residue 1-109 and V_H_ domain as residue 1-117. The elbow angles were calculated for the *P*1 and *P*6_5_ crystal structures, the free and binding states of the Certolizumab, as well as the trajectories from the molecular dynamics based on full-residue *P*1 and *P*6_5_ structures.

## 3. Results

### 3.1 Overall description of A33 Fab structure

A33 Fab is a humanised fragment consisting of a γ heavy chain and a κ light chain. Each chain contains a variable domain (V_L_ and V_H_) and a constant domain (C_L_ and C_H_1). The variable framework sequence is derived from the human antibody LAY and substituted with murine CDRs while the constant domain fully originated from mammal cells (King, Antoniw et al. 1995). The elbow angles, defined as the intersection angle of the two pseudo-dyad axes (PDAs) between the variable domain and the variable domain, of structures solved in *P*1 and *P*6_5_ space groups are 156° and 145° respectively, which implies flexibility of the switch region (Stanfield, Zemla et al. 2006).

A33 Fab illustrates a canonical β-sandwich Ig fold within four domains (V_L_, V_H_, C_L_ and C_H_1). Each domain has two layers of β-sheets, an inner- and an outer β-sheet. One canonical disulphide bridge was identified in each domain between the β-sheets layers (between _heavy_C22-_heavy_C96 in V_H, light_C_23-light_C_88_ in V_L, heavy_C_144-heavy_C_200_ in C_H_ and _light_C_134-light_C_194_ in C_L_), which contributes to the stability of the Fab (Figure 1). An inter-chain disulphide bridge, _light_C_214-light_C_220_, was not resolved due to the missing loop in the hinge region of the heavy chain. Seven inter-chain hydrogen bonds have been identified between the heavy and light chains, which are _heavy_P102-_light_Y36, _heavy_F103-_light_T46, _heavy_W106-_light_T46, _heavy_Q39-_light_Q38, _heavy_Q39-_light_Q38, _heavy_P170-_light_S162 and _heavy_P126-_light_S121. In addition to hydrophobic interactions, there are five inter-chain hydrogen bonds in variable regions while only two in constant domains, illustrating tighter contacts within variable regions.

**Figure 1.**
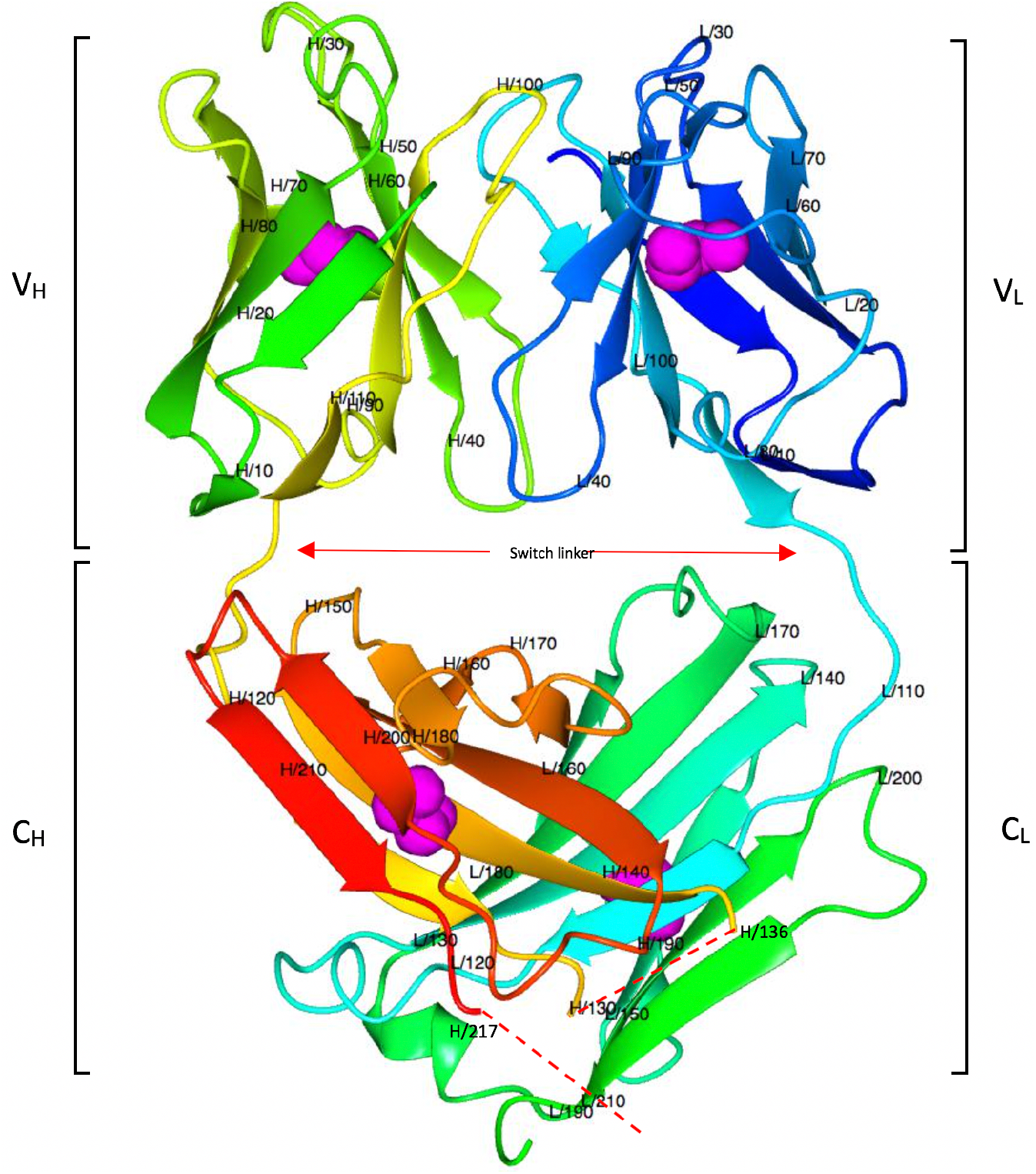
Overall A33 Fab structure. Residue numbers are labelled every ten amino acids. A33 Fab illustrates a canonical β-sandwich Ig fold within four domains (V_L_, V_H_, C_L_ and C_H_1) and each domain contains a disulphide bridge between the inner and outer β-sheets (magenta sphere). Missing loop regions are marked with a red dash.

### 3.2 Comparison of A33 Fab structures in different space groups

The triclinic crystals (*P1*) diffracted to 2.5 Å resolution with two copies of A33 Fab per asymmetric unit (AU), while the hexagonal crystals (*P6_5_*) have only one copy per AU and diffract to 2.2 Å. The overall structures in both crystal forms are similar but display slight differences (Figure 2A). The main difference between the two structures is the elbow angle. Different compact patterns affect the intersection angle between the variable region and the constant region. The elbow angles shift from 156° in the triclinic form to 145° in the hexagonal form, which implies flexibility of the switch region. However, the variable regions and the constant regions share great similarity. The RMSD calculated between Cα-atoms of matched residues at 3D superposition of the two structures is 1.33 Å. In contrast, the RMSD decrease to only 0.5 and 0.76 Å for separately superposed variable- and constant regions, respectively.

**Figure 2.**
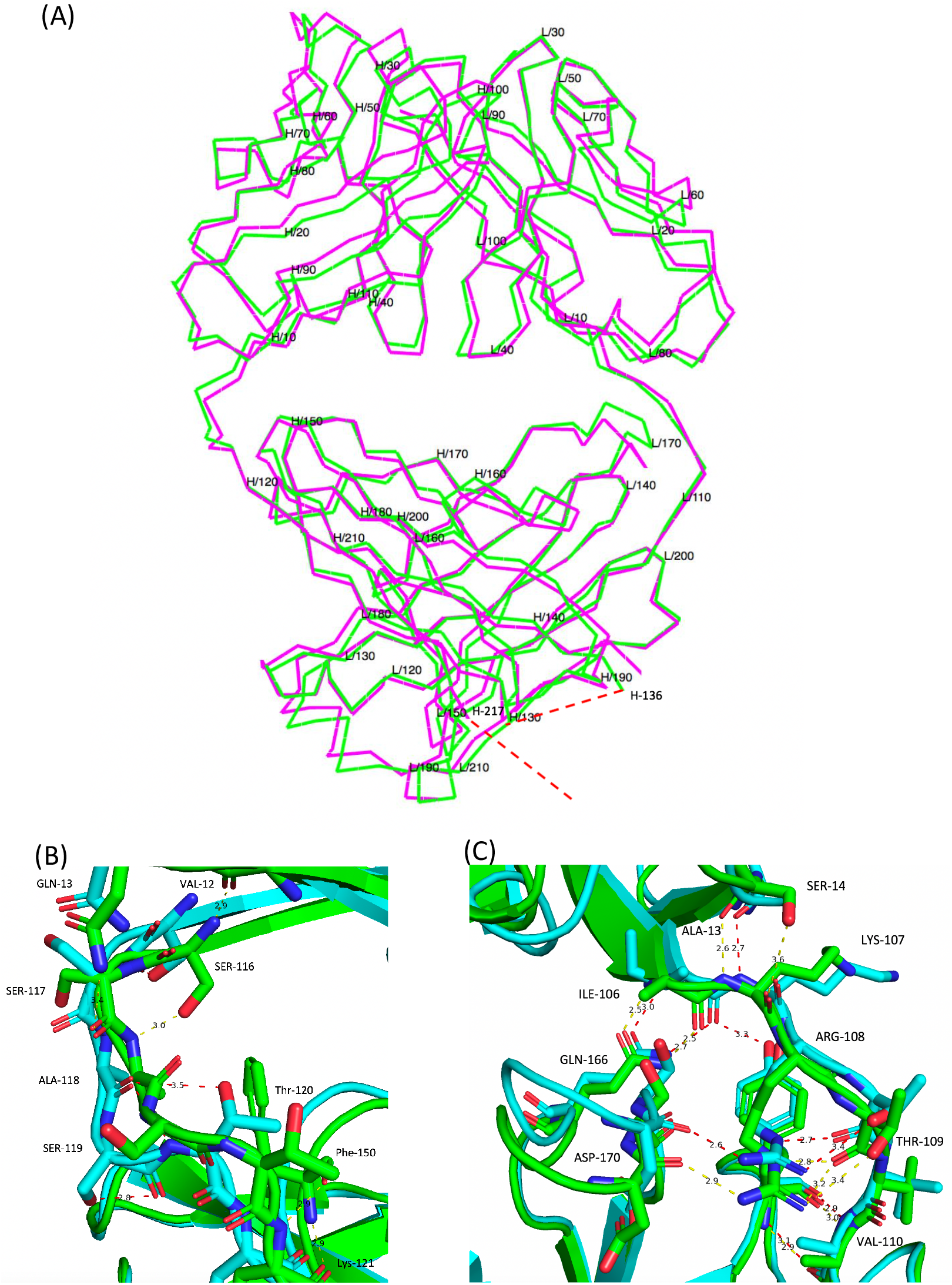
(A) Superposed C^α^ trace of A33 Fab structures in space groups *P*1 (magenta) and *P*6_5_ (green). The missing regions (_heavy_S132 to _heavy_T135 and _heavy_K218 to _heavy_A228) are marked with red dash lines. Residues are labelled every 10 amino acids. The two structures share high similarity but display differences in the switch region. (B) Superposed heavy chain switch region. The triclinic model is coloured in green while the hexagonal model is coloured in cyan. Conformational differences can be viewed from _heavy_S117 to _heavy_S119 while light chain switch peptides are fixed by a set of hydrogen bonds. (C) Superposed light chain switch region.

Closer insight into the linkers between the variable and the constant regions revealed conformational differences between the two models in heavy chain switch regions. The side chains of _heavy_S117 and _heavy_S119 have different orientations and form different hydrogen bonds with adjacent residues. Also, two unique hydrogen bonds are formed between _heavy_A118 and adjacent residues in the hexagonal model. In contrast, the linker in the light chain is fixed by a set of hydrogen bonds, which grants it less flexibility (Figure 2B, 2C).

Two loops are missing in both structural models, which indicates high flexibility in this region. In the C_H_ domain the missing loop is composed of four residues (SKST, _heavy_S132 to _heavy_T135) in the hexagonal model and six residues (SKSTSG, _heavy_S132 to _heavy_G137) in the triclinic model. The hydrophilicity of the missing loop could facilitate interactions with solvents and thus, contribute to the flexibility of the loop. The other missing loop is located at the C-terminus of the heavy chain, which is also the hinge region of A33 Fab. The missing loop in this region contains 11 residues (KSCDKTHTSAA, _heavy_K218 to _heavy_A228).

### 3.3 Elbow angles for the *P*1 and *P*6_5_ structures obtained from simulations

As shown in Figure 3 A, B, the elbow angles in the *P*1 and *P*6_5_ structures were initialised at 151° and 148.7°, respectively. In the *P*1 structure, this angle remained at around 150° for the first 15 ns before dropping to 144° at 17 ns onwards. The *P*6_5_ structure briefly dipped to 138° at 6 ns, but returned to 145° from 15 ns onwards. From 20 ns onwards, the elbow angles in both the *P*1 and *P*6_5_ structures fluctuated slightly in the range between 137° to 146°. This result implies that at these simulation conditions (pH 7.0), the elbow-angle in both structures rapidly equilibrated and converged on the elbow angle observed in the *P*6_5_ crystal form. Figure 3 shows the distribution of elbow angles for the two structures throughout their respective 100 ns simulations. Compared to the P65 structure, the *P*1 structure had a marginally higher frequency at 135-140° and at >150°. Nevertheless, the difference between the two structures was negligible.

**Figure 3.**
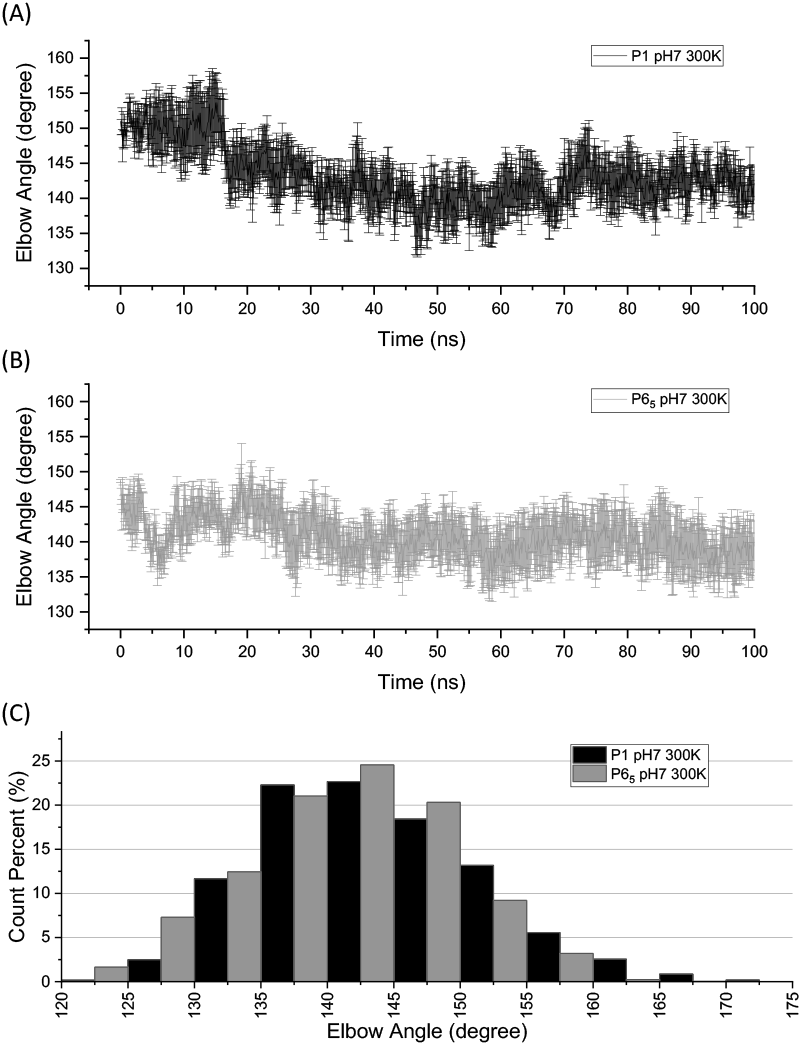
The elbow angles for the *P1* and *P6_5_* structures obtained from simulations. The average elbow angles were shown for the *P*1 (A) and *P*65 (B) structures during the 100 ns simulation at pH 7.0 and 300 K. The error bars represent standard errors of the mean (SEM). (C) Histogram for the elbow angle distribution in the *P*1 and *P*65 trajectories obtained during the simulations. In all figures, *P*1 and *P*65 are coloured in black and grey, respectively.

In summary, the elbow angle simulation implies that the *P*6_5_ structure more closely resembles the average conformation sampled in the simulation, compared to the *P*1 structure, and therefore better represents the structure of A33 Fab likely to be observed in solution.

### 3.4 Humanised complementarity determining regions (CDRs)

A33 Fab contains a highly humanised variable region composed of human variable framework regions (FWRs) from the LAY antibody and murine CDRs to minimise the formation of human anti-chimeric antibodies (HACAs)(King, Antoniw et al. 1995).

Here we aligned the A33 Fab for comparison with the peer therapeutic antibody Certolizumab which targets a different antigen (Lee, Shin et al. 2017). Certolizumab and A33 Fab have been developed from the same humanization scaffold with high sequence similarity (SI). The constant regions of A33 Fab and Certolizumab share high similarity. The RMSDs for C_H_ and C_L_ regions between A33 Fab and Certolizumab are 0.65 and 0.49 Å, respectively. By aligning the paratope regions, major differences can be found in CDR2 in V_L_ domains, CDR3 and CDR2 in V_H_ domains. Despite the large variation between CDRs, the FWRs remain highly similar (Figure 4B), which implies the rigidity of the FWRs.

**Figure 4.**
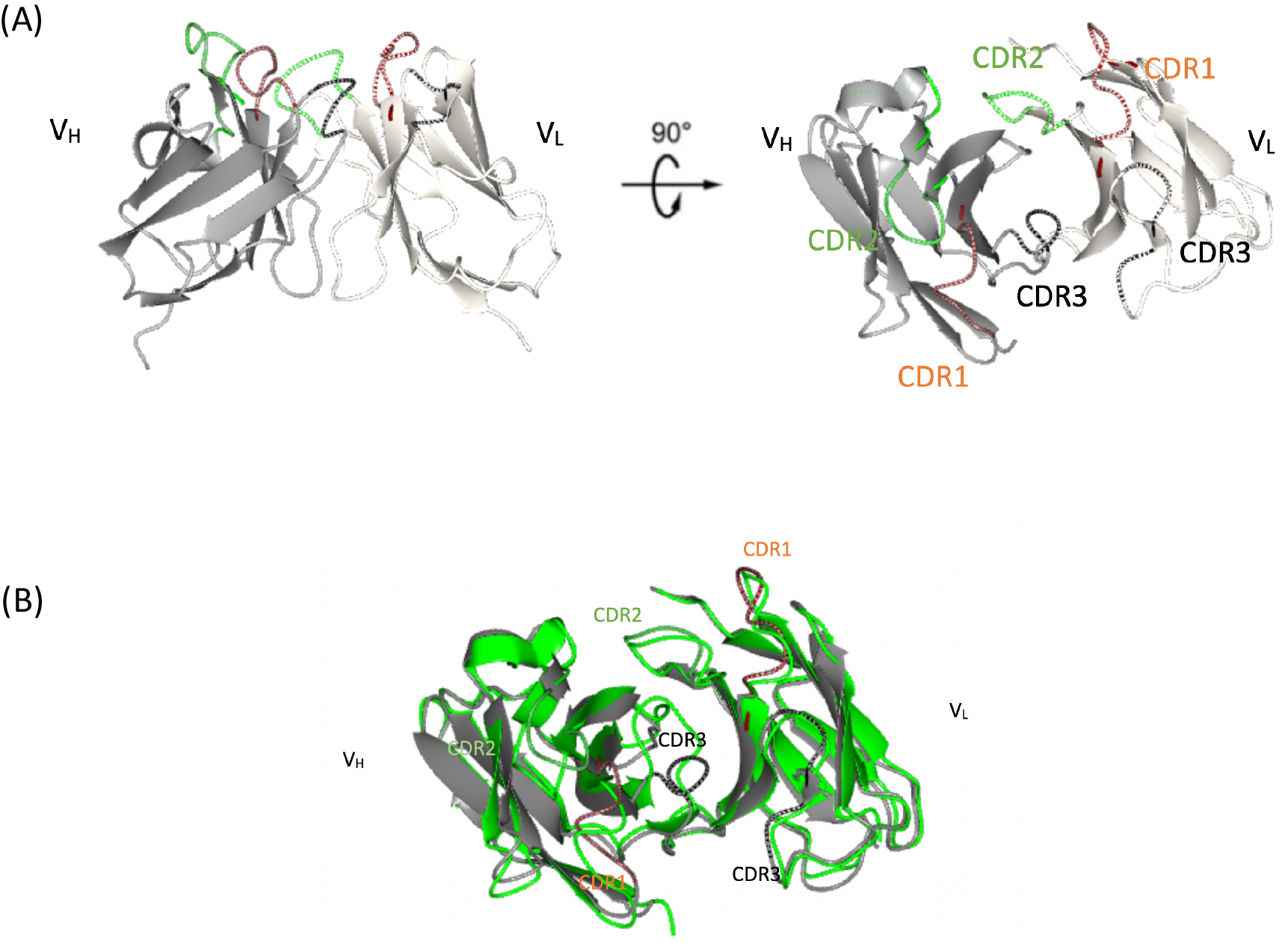
CDRs of A33 Fab. (A) CDR1 (brown), CDR2 (green) and CDR3 (black) originating from monoclonal murine A33 Fab are responsible for antigen binding. (B) Superposition of A33 FAB (grey) with Certolizumab (green).

### 3.5 Correlation between ΔΔ*G* and the *T*_m_, *ln*(*v*) based on crystal structures

To evaluate to what extent the crystal structures in *P1* and *P6_5_* could reflect the *in vitro* protein dynamics and stability, the *in silico* ΔΔ*G* was calculated for both structures for the mutations previously studied *in vitro* (Zhang, Samad et al. 2018). Due to the unresolved residues in the most flexible regions, the ΔΔ *G* was only calculated for eight and nine mutations in structures *P1* and *P65*, respectively. Figure ***5*.** (A) shows that the *P1* structure reveals a fairly good estimation of the protein’s thermal stability, *T*_m_, with R^2^ = 0.924. The correlation drops to 0.620 for aggregation kinetics *ln*(*v*), which is due to the additional involvement of colloidal stability in the aggregation that is not covered by ΔΔ*G*. For the ΔΔ*G* predictions based on the *P6_5_* structure, the correlation with *T*_m_ has a R^2^ of 0.681. Variants L/G66P and H/G178P deviate away from the linear regression line and result in decreased correlation compared to the *P1* structure (5**B**). In general, the correlations depend heavily on the destabilising variants and more unstable variants can be experimentally measured to evaluate the validity of the structure.

**Figure 5.**
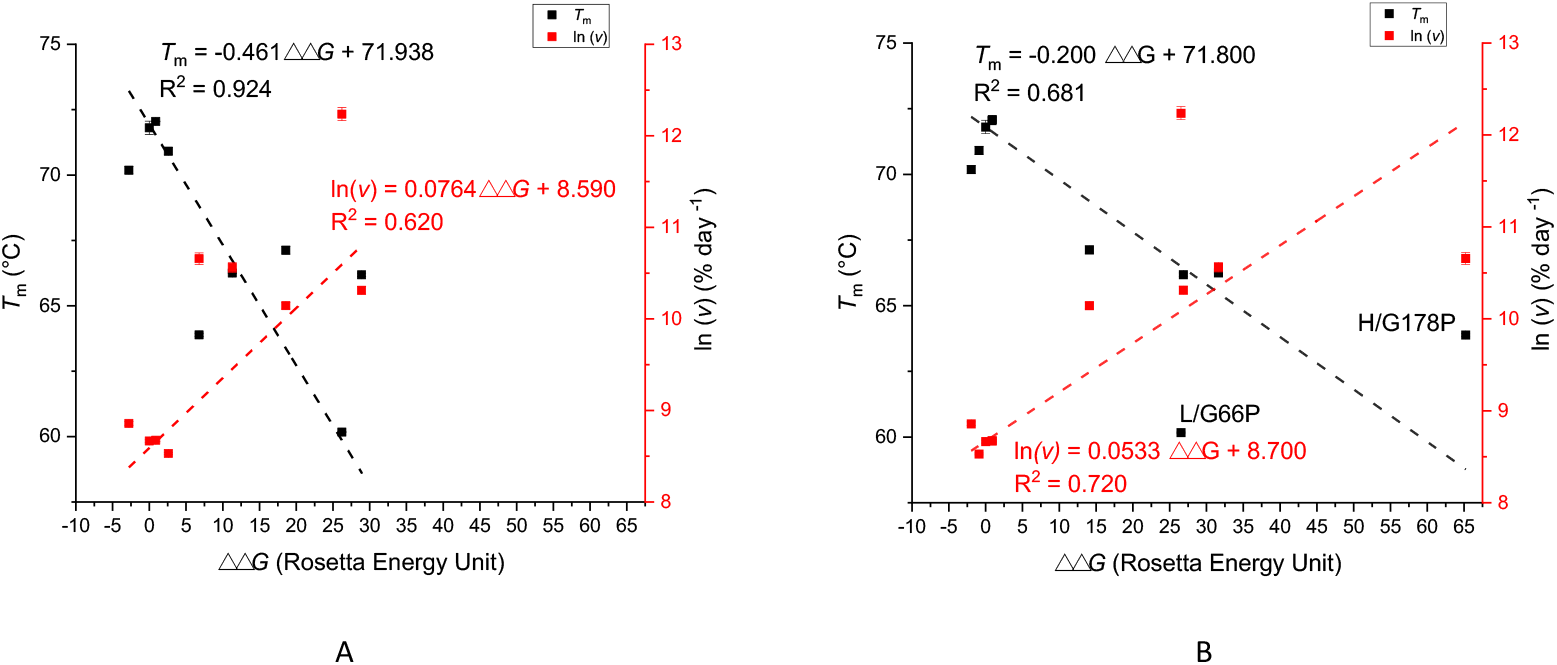
The correlation between ΔΔ *G* and *T*_m_ and aggregation kinetics *ln*(*v*) for the crystal structures obtained in *P_1_* (A) and *P6_5_* (B). ΔΔ*G* was calculated by Rosetta cartesian_ddg; *T*_m_ and *ln*(*v*) were measured previously (Zhang, Samad et al. 2018).

### 3.6 Correlation between ΔΔ*G* and the *T*m, *ln*(*v*) based on homology modelling from crystal structures

When flexible regions, such as unstructured loops, cannot be resolved by x-ray crystallography, an alternative is to conduct homology modelling to predict the missing residue conformations using protein modelling software (Kelley, Mezulis et al. 2015, Yang, Yan et al. 2015, Webb and Sali 2016). To evaluate the modelling accuracy, the predicted conformations need to be verified by *in vitro* data. In our work, *RosettaCM* was applied to construct the homology models, and Rosetta *cartesian_ddg* was used to predict the stability change upon point mutations. As Fab A33 shows high sequence identity (EMBOSS Needle) (Madeira, Park et al. 2019) of 90.1% and 88.3% to Certolizumab Fab (PDB entry 5WUV) and a human germline antibody (PDB entry 4KMT), respectively, the same procedure was also carried out with those PDB structures to compare the different Fab molecules (The homology model based on 4KMT was previously built by *minirosetta (Zhang, Samad et al. 2018)).*

As shown in the supplementary information (SI), the ΔΔ*G* correlation with *T_m_* and *ln*(*v*)depends largely on the five destabilising variants and the outliers for the prediction in the stabilising mutation. The correlation with *T*_m_ generally yields better results than that with *ln*(*v*). This is not surprising, as ΔΔ*G* could not cover colloidal stability on the protein surface. It is noticed that the H/Ser134Pro mutation had been predicted as an outlier in the models based on P1 (SI 2A), P65 (Version 1) (SI 2C) and Certolizumab Fab (SI 2G). In addition, H/Thr135Tyr arises as another outlier in P65 (version 1) (SI 2C), which decreases the correlation to an insignificant level (R^2^=0.033). A helix was predicted for residues _heavy_S132 to _heavy_T135 for P6_5_ (version 1) while this region was a loop in the other structures. As residues _heavy_S132 to _heavy_G137 represent one of the most flexible regions in the Fab structure, it is unlikely to form a structural helix, though this PDB ranks the top after homology modelling. As a result, a second top structure, P6_5_ (version 2) (**Fehler! Verweisquelle konnte nicht gefunden werden.**E, 2F), was selected with a loop predicted in that region. This selection eliminates all outliers, including H/Ser134Pro, around that loop. The model based on 4KMT also does not contain outliers in the stable mutations. By zooming in at the region _heavy_S132 to _heavy_G137, the loop conformations without outliers (SI 3B, 3D) both have the loop stretching outward at _heavy_S132 position, which could possibly avoid high energy penalties caused by the H/Ser134Pro mutation.

The hinge conformation is also not consistent in different batches of homology modelling. Though the model based on 4KMT yields superior correlation, it is unlikely for this flexible region to be helical at the hinge tail (SI 2J). To examine the possible conformations for hinge regions, the homology models were further studied using molecular dynamics. Six repeats were performed for each model. However, after 100 ns, none of the hinge regions in the trajectories converged to a relatively consistent conformation (SI). Therefore, the hinge conformation is very versatile and undoubtedly a challenge to be resolved. Multiple stabilising mutations at or near the hinge region could be introduced to rigidify the flexibility, increasing the chances to reduce this flexibility.

The three structures that yielded the best correlations with *T*_m_ and *ln*(*v*), namely the models based on P1, 5WUV and 4KMT, are compared directly for their loop (residues H/132-137, **Fehler! Verweisquelle konnte nicht gefunden werden.**P) and hinge (residues H/218-228 (**Fehler! Verweisquelle konnte nicht gefunden werden.**R) regions, and with their RMSD calculated in reference to structure P1 (Table 2). The model based on 4KMT yields a higher RMSD (0.337 nm) compared to the one based on 5WUV (0.311 nm), probably owing to the 2% lower sequence identity in 4KMT. To investigate the local displacement of the predicted loop and hinge, these three structures are also aligned based on the loops or their adjacent beta-strands, (SI 2L, 2N). This quantifies the relative contributions from different positions and forms of the loop. The results show that the hinge conformations yield much higher RMSD, especially when the RMSD is calculated based on the alignment of its adjacent β-strands. This further verifies the versatility and flexibility of hinge conformations.

**Table 1.**
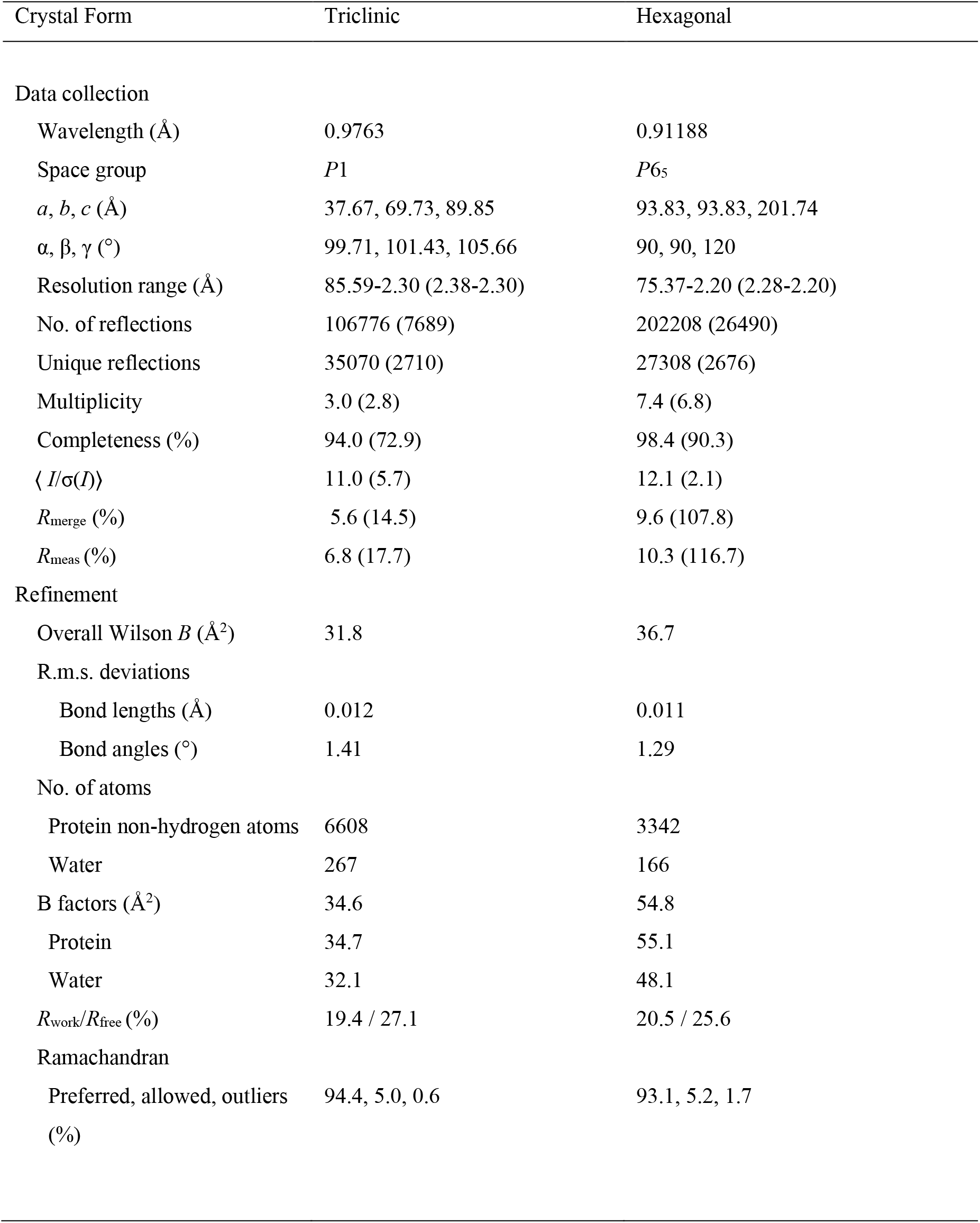
Data collection and structure refinement statistics for A33 FAB in space groups P1 and P65. Values for the outer shell are given in parentheses.

**Table 2.**
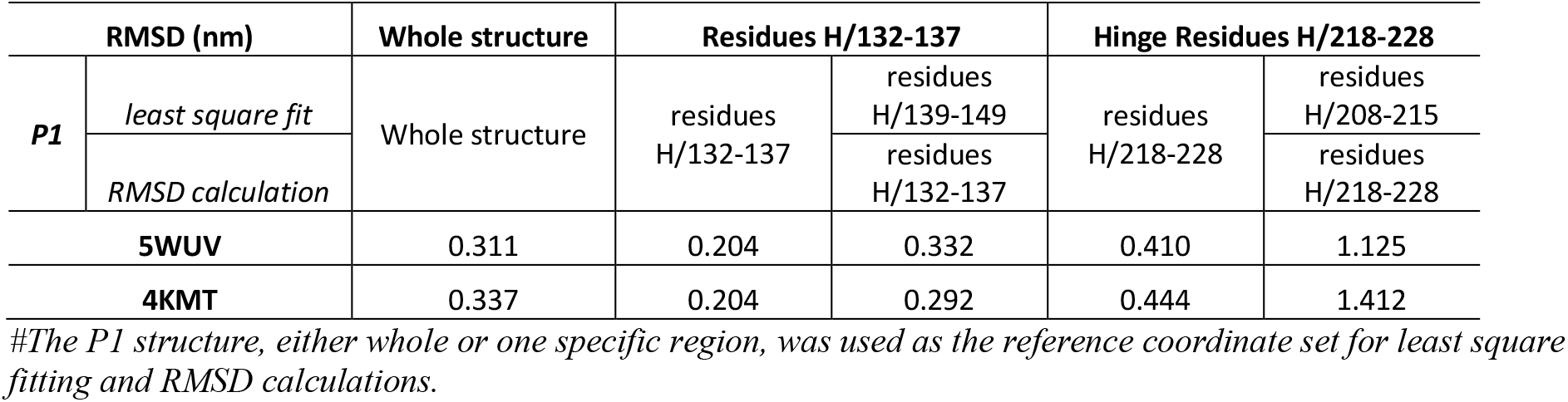
The RMSD between the structures with the highest correlation with *in vitro* data.

## 4. Discussion

Over the past decades, significant efforts have been undertaken into the development of therapeutic antibodies against CRC targeting human antigen A33. Various A33 Fab variants have been developed and multiple clinical trials have been carried out (Welt, Ritter et al. 2003, O’Donoghue, Smith-Jones et al. 2011). However, various A33 Fab variants have failed in clinical trials due to the normal gut localization and intrinsic stability issues of the variants (O’Donoghue, Smith-Jones et al. 2011, Codina Castillo 2019). To our knowledge, the structure of A33 Fab or its variants has never been reported. Here we determined the structure of a A33 Fab mutant, H/C226S, in two crystal forms.

The structural analysis of A33 Fab in two different crystal forms disclosed the intrinsic rigidity of four separate regions (V_L_, C_L_, C_H1_, C_L_) and the flexibility of the switch region. The different compact patterns and the interactions with solvents altered the elbow angles between the two pseudo-dyad axes from 156° (triclinic form) to 145° (hexagonal form), which leads to the noticeable RMSD of the structures between the two crystal forms. In contrast, the canonical β-sandwich Ig folds contribute to the rigidity of both variable and constant regions, which leads to the high similarity of separate domains in different crystal forms. Subsequent simulations illustrated that the elbow angles of both structures are prone to equilibrate and converge on an average of 145°, which implied that the *P*6_5_ structure is the most likely representation of the average solution structure of A33 Fab at pH 7.0.

To investigate the similarity of the elbow angle connecting variable and constant domains, molecular dynamics were performed on the full-residue homology models of *P1*, *P6_5_* and Certolizumab. Principle Component Analysis (PCA) was performed to capture the major collective motions throughout the simulation (SI). The PC1 motion illustrates the elbow bending, whereas all the three structures share overlapped distribution when their trajectories are projected onto the PC1 eigenvectors. This implies that the three structures, though having minor sequence difference, would undergo resembled dynamics in the elbow bending movement.

We hypothesised that the major differences between the A33 Fab and Certolizumab are located at the variable region. Certolizumab has been developed from the same humanization scaffold as A33 Fab, possessing a similar variable region framework and the same constant region protein sequence. To our surprise, the variable regions of these two Fabs share higher similarities than expected (RMSD 0.91 Å). Only the V_H_ CDR3 and V_H_ CDR2 of both Fabs vary a lot while the rest of the CDRs have similar conformations even with different protein sequences, which suggests the key role of V_H_ CDR3 and V_H_ CDR2 in the identification between human A33 and TNFα. The FWRs are also similar between the two structures despite the sequence differences, which implies that the FWRs offer good support to the CDRs while being neutral to scaffolding for antibody structural integrity. The other main difference is the elbow angle. According to the previously published Certolizumab structure, the elbow angle of Certolizumab changed 9° during antigen binding, from 138° (free) to 129° (binding) (Lee, Shin et al. 2017). In contrast, the elbow angles of *P*1 and *P*6_5_ structures of A33 Fab are 156° and 145° respectively. Certolizumab and Fab A33 possess the same protein sequence in switch and constant regions while the elbow angles differ, which implies flexibility in the switch region.

It has been reported that the flexibility of elbow angle is vital for the binding affinity of Fabs to antigens by optimizing antibody conformational dynamics and adaptation to antigen structure (Lord, Bird et al. 2018, Henderson, Watts et al. 2019). In conclusion, the scaffold of the A33 Fab is optimal to support CDRs targeting different antigens and has the potency to be applied to other therapeutic candidates from an engineering perspective.

The two structures could not resolve two flexible regions, including residues _heavy_S132 to _heavy_T135 and _heavy_K218 to _heavy_A228. Thus, homology modelling was conducted to predict the missing residues, and *in vitro T*_m_ and aggregation kinetics *ln*(*v*) were used to validate the prediction accuracy. *In silico* ΔΔ*G* correlation illustrated that the *P1* structure reveals better estimation on the protein thermal stability. As the correlation is based on the mutants previously studied *in vitro*, it implies that the *P1* model could reveal a natural conformation *in vitro* while the *P6_5_* model discloses a slighted altered conformation.

Further efforts were undertaken to predict the conformation of loop regions missing in the x-ray models. By using ΔΔ*G* correlations, we picked out the best solutions and aligned them with other models showing high homology. Unsurprisingly, the predicted loop regions illustrate low similarity between each other. In *P1* and *P6_5_* models, residues H/132-142 comprise loops with different conformations while residues H/217-228 form hinge tails pointing at completely different orientations. The comparison between *P1* and other homology models further validated the versatility and flexibility of these loops, especially at the hinge region located at the C terminal of the proteins.

For the first time, the conformation of CDRs and the humanization framework of A33 Fab are reported, which may provide guidance to the generation of novel A33 Fab mutants with higher affinity and less immunogenicity. The structural comparison of A33 Fab with Certolizumab provides a novel perspective of how humanised Fabs developed from the same human Ig framework and differ from each other in terms of targeting different antigens. Based on the A33 Fab crystal structure we are carrying out stability studies by molecular dynamics simulation. Further structural work is needed to acquire the co-crystallization structure of A33 Fab to A33 antigen, which will disclose the interaction of the CDRs to the target and the conformational changes during binding.

## Supporting information

Supplementary material section for Fab manuscript

## Acknowledgements

We would like to thank Diamond Light Source for beamtime (proposal mx23853), and the staff of beamlines I04-1 and I24 for assistance during data collection. We are grateful for funding support to Jiazhi Tang from the Chinese Scholarship Council. We thank UCB Pharma for providing the A33 Fab. We also gratefully acknowledge the Engineering and Physical Sciences Research Council (EPSRC) for funding (EP/N025105/1) that supported Cheng Zhang.

