## Supplementary material section for Fab manuscript for "The structure of the humanised A33 Fab C226S variant, an immunotherapy candidate for colorectal cancer"


### Supplementary Information:

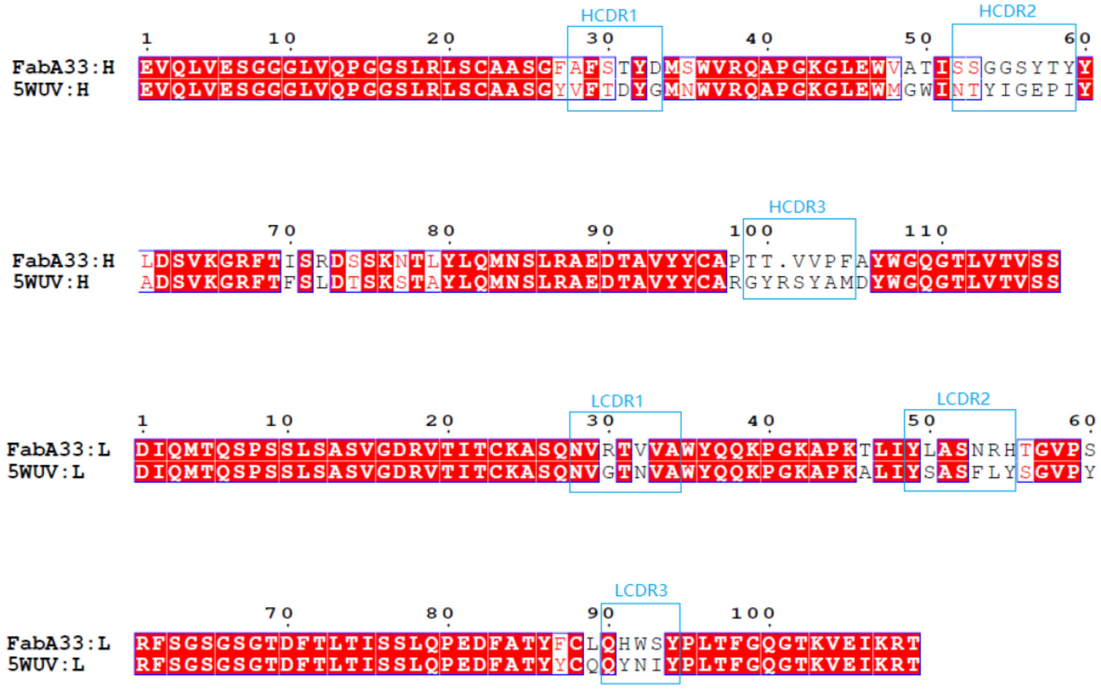


**Figure S1.** Alignment of the protein sequences of variable domains between A33 Fab and Certolizumab.

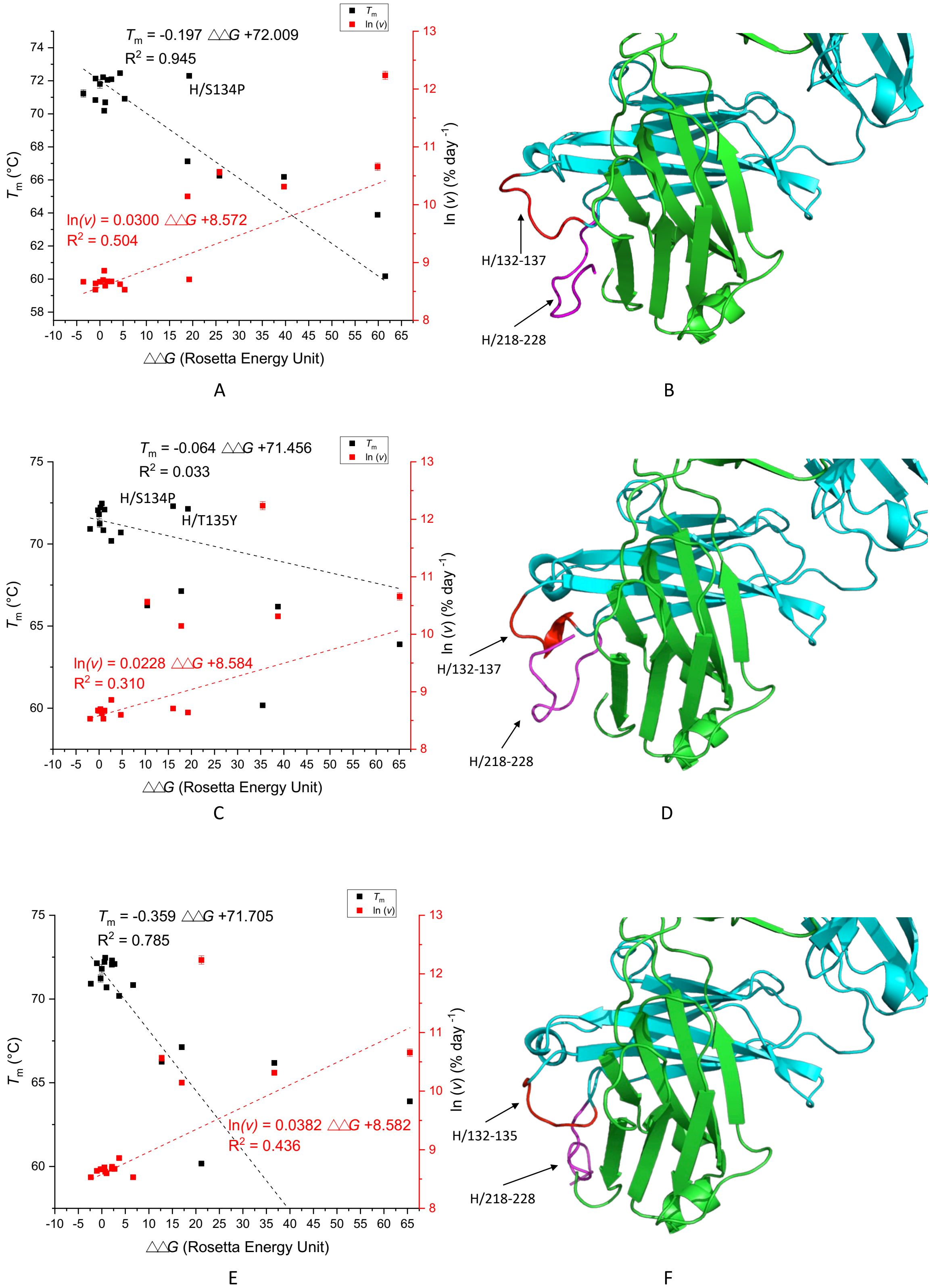

$T_m$ = -0.197 △△G +72.009
$R^2$ = 0.945
H/S134P
ln(v) = 0.0300 △△G +8.572
$R^2$ = 0.504
$T_m$
ln (v)
$T_m$ (°C)
ln (v) (% day$^{-1}$)
△△G (Rosetta Energy Unit)
A
H/132-137
H/218-228
B
$T_m$ = -0.064 △△G +71.456
$R^2$ = 0.033
H/S134P
H/T135Y
ln(v) = 0.0228 △△G +8.584
$R^2$ = 0.310
$T_m$
ln (v)
$T_m$ (°C)
ln (v) (% day$^{-1}$)
△△G (Rosetta Energy Unit)
C
H/132-137
H/218-228
D
$T_m$ = -0.359 △△G +71.705
$R^2$ = 0.785
ln(v) = 0.0382 △△G +8.582
$R^2$ = 0.436
$T_m$
ln (v)
$T_m$ (°C)
ln (v) (% day$^{-1}$)
△△G (Rosetta Energy Unit)
E
H/132-135
H/218-228
F

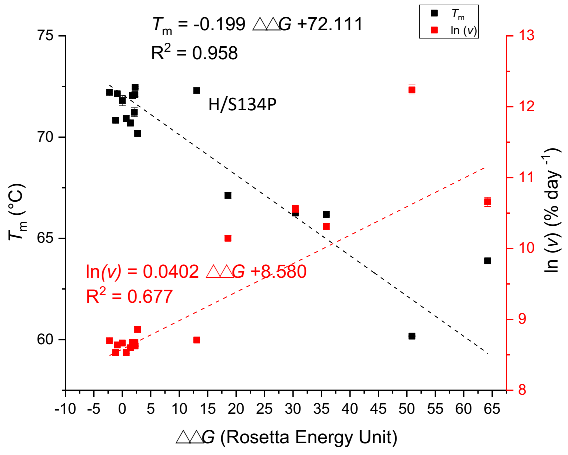
$T_m$ = -0.199 △△G +72.111
$R^2$ = 0.958
$T_m$
ln (v)
H/S134P
ln(v) = 0.0402 △△G +8.580
$R^2$ = 0.677
$T_m$ (°C)
ln (v) (% day -1)
△△G (Rosetta Energy Unit)
75
70
65
60
13
12
11
10
9
8
-10 -5 0 5 10 15 20 25 30 35 40 45 50 55 60 65

G

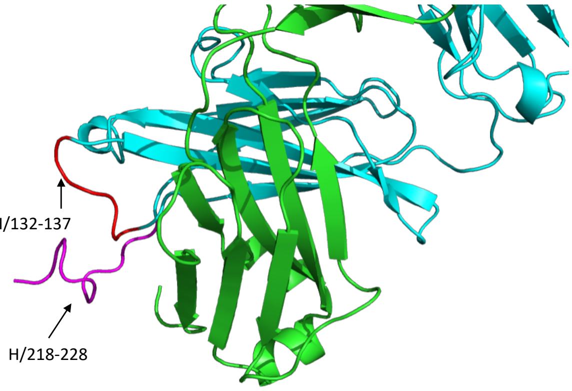
H/132-137
H/218-228

H

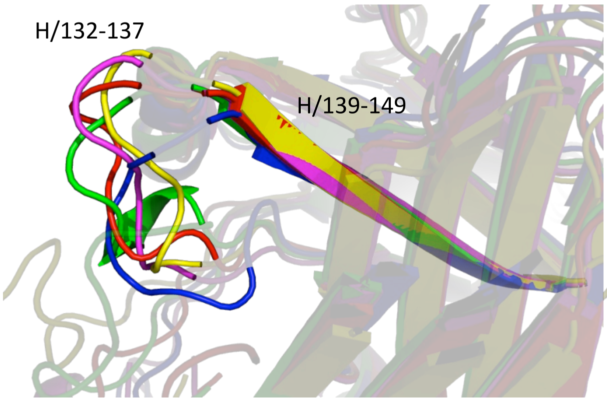
H/132-137
H/139-149

K

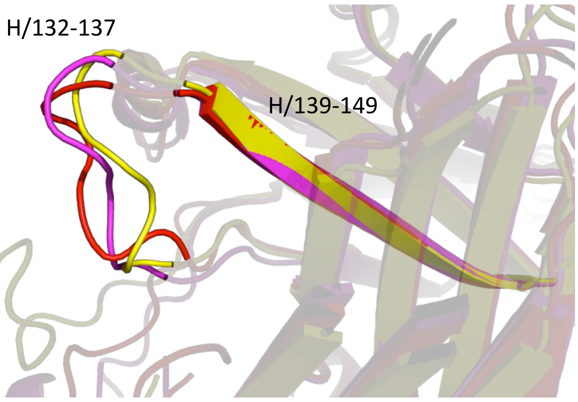
H/132-137
H/139-149

L

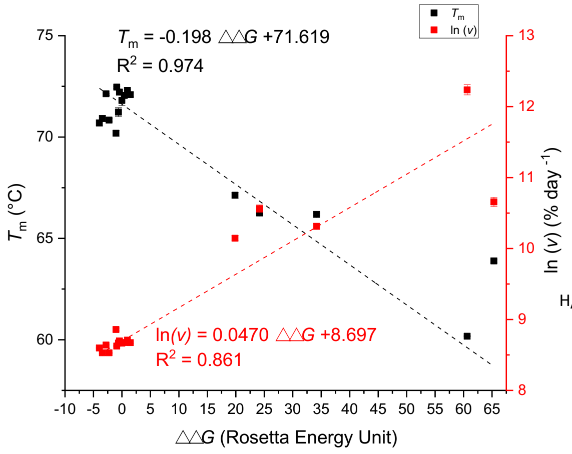
$T_m$ = -0.198 △△G +71.619
$R^2$ = 0.974
$T_m$
ln (v)
ln(v) = 0.0470 △△G +8.697
$R^2$ = 0.861
$T_m$ (°C)
ln (v) (% day -1)
△△G (Rosetta Energy Unit)
75
70
65
60
13
12
11
10
9
8
-10 -5 0 5 10 15 20 25 30 35 40 45 50 55 60 65

I

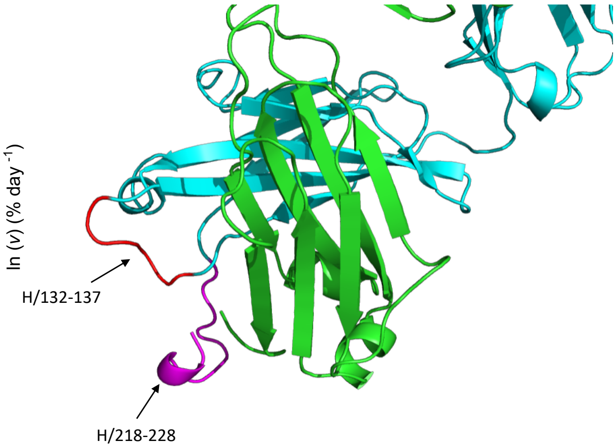
H/132-137
H/218-228

J

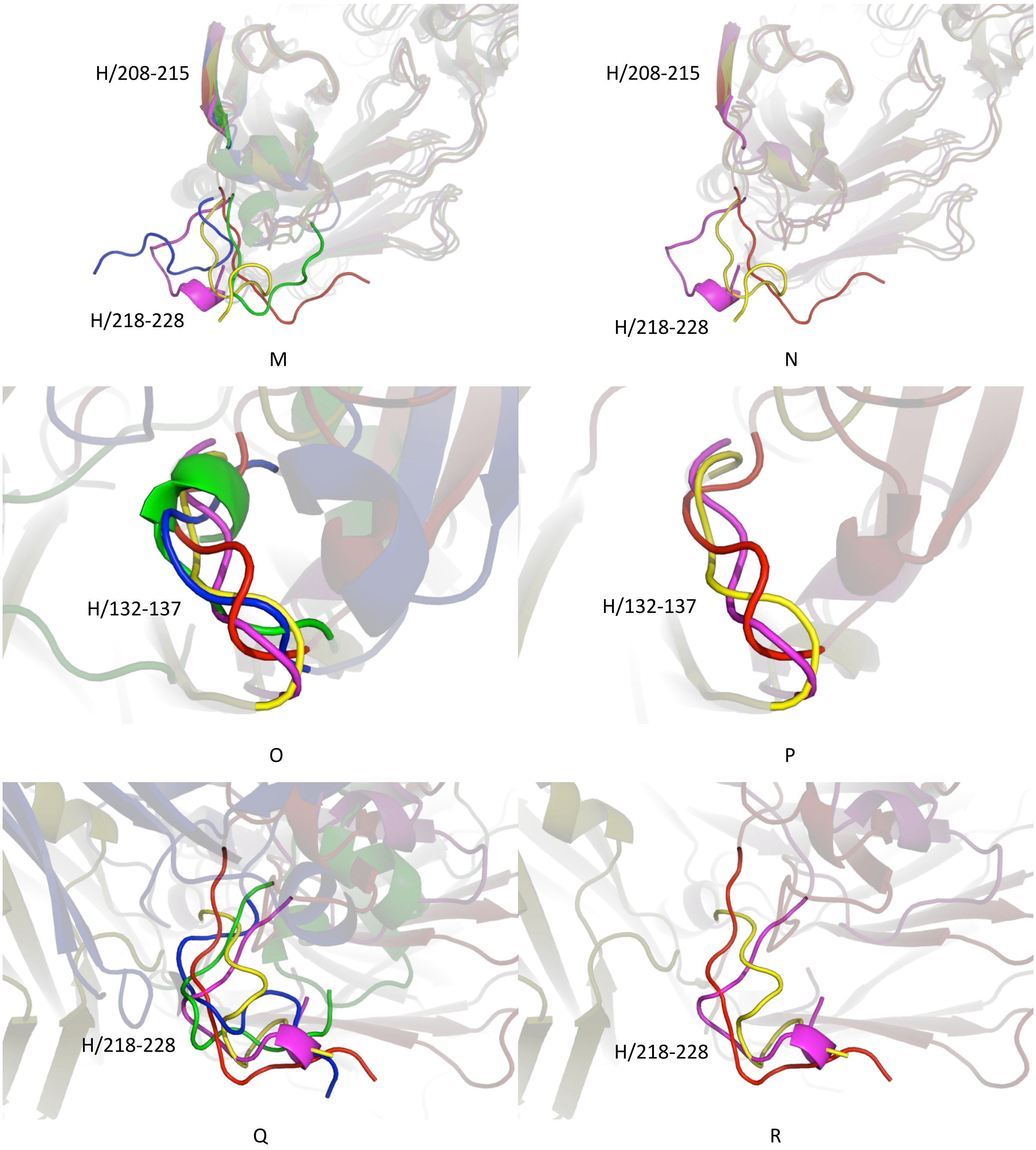


**Figure S2.** The correlation between ΔΔ*G* and $T_m$ and aggregation kinetics *ln(v)* for the full-length homology models based on the crystal structures. Figure A, C, E, G and I are the correlations for homology modelling based on *P1*, $P6_5$ (version 1), $P6_5$ (version 2), Certolizumab Fab (PDB entry 5WUV) and a human germline antibody (PDB entry 4KMT), respectively. Figure B, D, F, H and J highlight the conformations prediction for residues H/132-137 (red) and residues H/218-228 (magenta) based on *P1*, $P6_5$ (version 1), $P6_5$ (version 2), Certolizumab Fab (5WUV) and a human germline antibody (PDB entry 4KMT), respectively. Light and heavy chains are coloured in green and cyan, respectively. The homology modelling based on *P1* (red), $P6_5$ (Version 1) (green), $P6_5$ (Version 2) (blue), 5WUV (yellow) and 4KMT (magenta) are locally aligned to β-strand residues H/139-149 in SI 2K or aligned to β-strand residues H/208-215 in SI 2M; SI 2L and 2N are the subsets of SI 2K and 2M, respectively, only showing the models based on *P1* (red), 5WUV (yellow) and 4KMT (magenta). The five homology models, using the same colouring scheme, are also locally aligned to residues H/132-137 (SI 2O) or residues H/218-228 (SI 2Q), where SI 2P and 2R only show the *P1*, 5WUV and 4KMT.

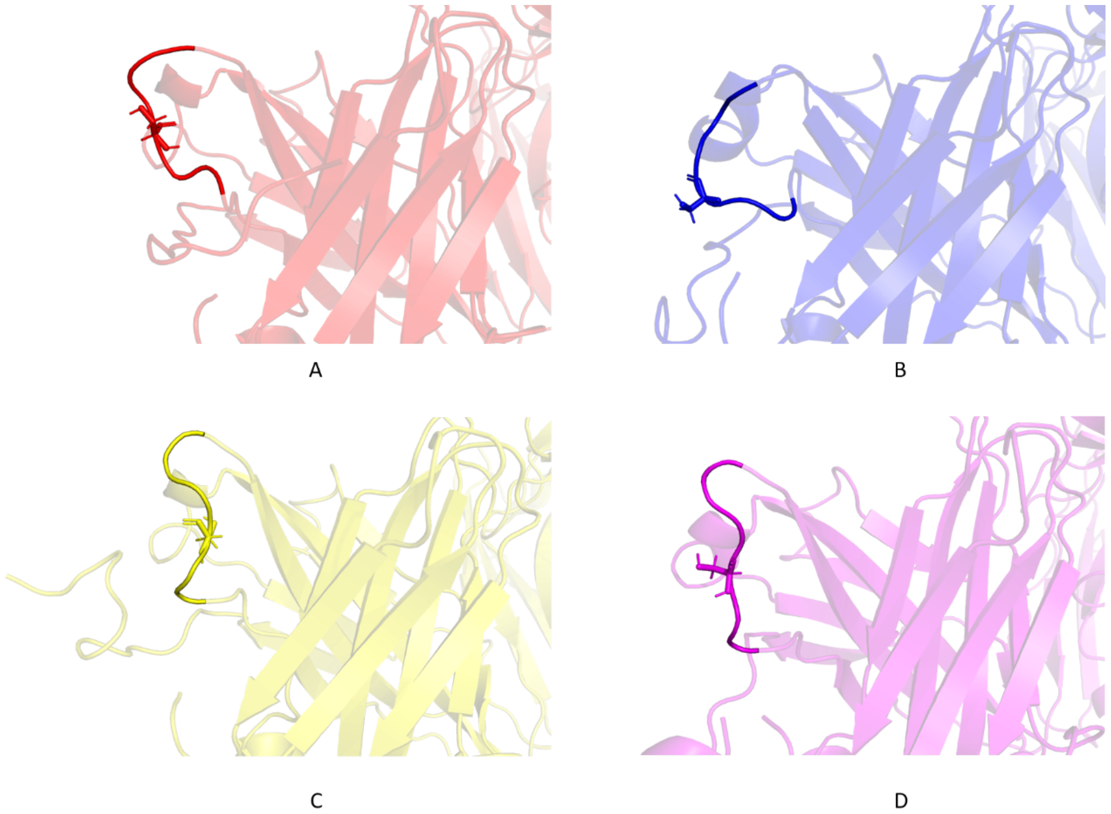


**Figure S3.** The loop conformation for residues H/132-137 of the homology models. The remaining part of the structure is partially transparent with residue H/Ser134 shown in sticks. Homology models are built based on *P1* (A), $P6_5$ (Version 2) (B), Certolizumab Fab (PDB entry 5WUV) (C) and a human germline antibody (PDB entry 4KMT).
